# Effects of dopamine agonist treatment on resting-state network connectivity in Parkinson’s disease

**DOI:** 10.1101/2020.01.31.928853

**Authors:** David M. Cole, Bahram Mohammadi, Maria Milenkova, Katja Kollewe, Christoph Schrader, Amir Samii, Reinhard Dengler, Thomas F. Münte, Christian F. Beckmann

## Abstract

Dopamine agonist (DA) medications commonly used to treat, or ‘normalise’, motor symptoms of Parkinson’s disease (PD) may lead to cognitive-neuropsychiatric side effects, such as increased impulsivity in decision-making. Subject-dependent variation in the neural response to dopamine modulation within cortico-basal ganglia circuitry is thought to play a key role in these latter, non-motor DA effects. This neuroimaging study combined resting-state functional magnetic resonance imaging (fMRI) with DA modification in patients with idiopathic PD, investigating whether brain ‘resting-state network’ (RSN) functional connectivity metrics identify disease-relevant effects of dopamine on systems-level neural processing. By comparing patients both ‘On’ and ‘Off’ their DA medications with age-matched, un-medicated healthy control subjects (HCs), we identified multiple non-normalising DA effects on frontal and basal ganglia RSN cortico-subcortical connectivity patterns in PD. Only a single isolated, potentially ‘normalising’, DA effect on RSN connectivity in sensori-motor systems was observed, within cerebro-cerebellar neurocircuitry. Impulsivity in reward-based decision-making was positively correlated with ventral striatal connectivity within basal ganglia circuitry in HCs, but not in PD patients. Overall, we provide brain systems-level evidence for anomalous DA effects in PD on large-scale networks supporting cognition and motivated behaviour. Moreover, findings suggest that dysfunctional striatal and basal ganglia signalling patterns in PD are compensated for by increased recruitment of other cortico-subcortical and cerebro-cerebellar systems.

## Introduction

Parkinson’s disease (PD) is a neurodegenerative disorder characterised by hypo-dopaminergic neurotransmission within the nigrostriatal dopamine pathway (Fahn, 2008), leading to pronounced motor deficits. In addition, many sufferers experience ‘higher-order’ cognitive and neuropsychiatric symptoms, such as impairments in working memory, decision-making and motivation (Owen et al. 1992; Aarsland et al. 1999; Dagher and Robbins, 2009; Gao and Wu, 2016). Growing evidence suggests that, in a notable minority of PD patients, such non-motor, higher-order symptoms may be related to the same dopamine agonist (DA) medications used to treat primary motor problems (Cools, 2006; Dagher and Robbins, 2009; Cilia and van Eimeren, 2011; Antonelli et al, 2014). In particular, higher-order cognitive side effects of DA treatment often present as increased impulsive and compulsive behaviours and, in severe cases, neuropsychiatric syndromes, such as addictions and impulse control disorders (Dagher and Robbins, 2009; Weintraub et al. 2010; Vargas and Cardoso, 2018).

It has been suggested that: “Persistent pharmacological stimulation of dopamine receptors [in] PD patients, could [result in] increased engagement in reward-seeking behaviours and reduced ability to disengage [despite] negative consequences” (Dagher and Robbins, 2009). This statement applies principally to receptors within the mesolimbic dopamine pathway, which – in contrast to nigrostriatal neuroreceptors – do not suffer primary degeneration in PD, but may instead be hyper-stimulated by DA administration (Riba et al. 2008; Dagher and Robbins, 2009; Weintraub et al. 2010). Thus, although serving to ‘normalise’ motor deficits (see also Williams et al. 2002), DA medications may also effectively promote impulsive/compulsive behaviours via incidental mechanisms of neuroplasticity. The neural bases of these ‘non-normalising’ DA effects, while incompletely characterised, are thought to involve the dopamine-rich ventral striatum and connected regions of the mesolimbic pathway (Riba et al. 2008; Weintraub et al. 2010; Voon et al. 2011; Ye et al. 2011). Such regions comprise large-scale, cortico-striato-thalamic circuitry implicated in higher-order cognitive processes supported by dopamine, relevant for impulsivity and disturbed in PD (Honey et al. 2003; Everitt and Robbins, 2005; Dagher and Robbins, 2009; Hacker et al. 2012; Agosta et al. 2014; Ruitenberg et al. 2018). However, susceptibility to non-normalising DA effects varies across individuals, ostensibly in line with neurochemical and cognitive phenotypes (e.g., Dagher and Robbins, 2009; Buckholtz et al. 2010; Housden et al. 2010; O’Sullivan et al. 2011; Vargas and Cardoso, 2018). Importantly, this cognitive-neuropsychiatric variability could reflect individual differences in baseline dopaminergic functioning, leading to differential sensitivity of neuronal systems to subsequent drug modulation: particularly within distinct striatal and other subcortical regions (Cools, 2006), but also in higher-level cortical areas (Cools and D’Esposito, 2011; Voon et al. 2011; Ye et al. 2011).

Recent functional magnetic resonance imaging (FMRI) research, employing analysis methods sensitive to the synchronicity of activity, or ‘functional connectivity’, within large-scale brain networks, has suggested that sources of clinical-pharmacological variability may be explained by *systems-level* variability measurable in cortico-subcortical signalling patterns relevant for both dopamine function (e.g., in healthy subjects: Nagano-Saito et al. 2008; Cole et al. 2012; 2013) and dysfunction in PD (Kwak et al. 2010; Hacker et al. 2012; Helmich et al. 2012; Ruitenberg et al. 2018). For example, previously we have demonstrated evidence of specific, large-scale, higher-order neural systems possessing *task-independent* functionality relevant for dopaminergic modulations, as measured by ‘resting-state’ network (RSN) functional connectivity (Cole et al. 2012; 2013). This approach may thus help to resolve methodological and conceptual ambiguities in the *task-based* FMRI and related neuroimaging literature regarding the systems-level effects of DA therapeutics in PD (e.g., Hacker et al. 2012; Antonelli et al. 2014).

In the current study, we extended network-sensitive analysis techniques, applied previously to task-independent pharmacological FMRI data from healthy young adults (Cole et al. 2013), to investigate DA treatment effects in PD on RSN functional connectivity within cortico-subcortical circuitry thought to be dysfunctional in the disease. To test the hypothesis that DA treatments exert both beneficial (normalising) and detrimental (non-normalising) influences, relative to healthy control subjects, on distinct processing systems (sensory-motor and cognitive/motivational, respectively), we investigated within-subject connectivity changes during DA withdrawal in PD. We focussed on networks involving frontal cortex, ventral striatum and/or other basal ganglia regions due to their relevance for neuropsychiatric dysfunction and mesolimbic dopamine neuromodulation in PD. We also examined associations between disease- and DA-related striatal functional connections and a behavioural measure of impulsive decision-making that quantified participants’ ‘delay discounting’ of monetary reward; the intertemporal choice (ITC) questionnaire (Kirby et al. 1999). As variation in both delay discounting and impulsivity more generally have been related to functional processes recruiting striatal dopamine (Dalley et al. 2007; Riba et al. 2008; Buckholtz et al. 2010; Cai et al. 2011; Kayser et al. 2012), it was hypothesised that these functional connections would be differentially associated, across groups and conditions, with ITC-derived individual differences in impulsive choice.

## Materials and Methods

### Participants and study design

Fifteen adults diagnosed with idiopathic PD (mean age = 60 ± 7.4 SD, range = 42-74 years) and 19 HCs matched for age (mean = 61 ± 7.0, range = 44-69) and years of education (PD mean = 14 ± 2.6 SD, range = 11-19; HCs mean = 15 ± 3.0 SD, range = 11.5-22) completed the study. All subjects were drawn from a sample described previously (Milenkova et al. 2011; see also *Supplementary Materials and Methods*). All patients were being treated with DA medications primarily targeting dopamine D2/D3 receptors (12 treated with 1.05-3.50 mg pramipexole, two with 4-15 mg ropinirole and one with 300 mg piribedil). Eight of the 15 patients received DA monotherapy, while seven also required concomitant levodopa (L-dopa) medication (150-850 mg) that was not withdrawn during the study (see Discussion, *Methodological considerations*).

Patients underwent resting-state FMRI on two separate visits, two weeks apart and always in the morning. The first scan was conducted with patients in their normal medicated state (‘PD On’), while the second was conducted approximately 12 hrs after DA withdrawal (‘PD Off’). HCs were scanned only once, without any concomitant pharmacological challenge. Informed consent was obtained from all subjects and all study procedures were approved by the University of Magdeburg ethical committee and carried out in accordance with the standards of the Declaration of Helsinki.

### Intertemporal choice (ITC) questionnaire

The ITC questionnaire was administered at each visit, as described in detail in a recent publication examining behavioural data from subjects also included in the current FMRI study (Milenkova et al. 2011; see also *Supplementary Materials and Methods*).

### Image acquisition

Blood oxygenation level-dependent (BOLD) FMRI data were acquired for six minutes in the resting state on a 3T Siemens Magnetom Allegra MRI scanner. Participants were instructed to close their eyes during scanning. In total, 178 volumes of T2*-weighted echo planar imaging (EPI) data were acquired per session (TR = 2.0 s, TE = 30 ms, flip angle = 80º, voxel size = 3.0 × 3.0 × 3.0 mm, 34 slices with 0.75 mm gaps), angled parallel to the anterior-posterior commissural plane. A high-resolution (1 mm^3^ isotropic resolution) T1-weighted MPRAGE structural image was also acquired for each subject.

### Image preprocessing and network definition

Resting-state FMRI data were preprocessed with tools from the FMRIB Software Library (FSL; www.fmrib.ox.ac.uk/fsl; Smith et al. 2004) as described previously (Cole et al. 2013; see also *Supplementary Materials and Methods*). Preprocessing of each EPI dataset included removal of the first four volumes to allow for magnetic equilibration, motion correction, brain extraction, spatial smoothing with a Gaussian kernel of 5 mm FWHM, high-pass temporal filtering at 100 s and nonlinear registration to the MNI152 stereotactic template following affine registration to the associated high-resolution T1 structural scan.

To quantify differences in RSN cortico-subcortical functional connectivity between PD patients and HCs, as well as between On and Off DA treatment conditions in PD, seed-based partial correlation analysis (SBCA) was conducted. This approach measures the strength of BOLD signal correlation between voxels in a ‘seed’ region and those in a ‘target’ region or set of regions forming a network (O’Reilly et al. 2010; Cole et al. 2013). To test for PD and DA treatment-related effects on cortico-subcortical systems, we first created anatomically-defined subcortical seed masks for each subject (see *Supplementary Materials and Methods*). We next defined neural functional connectivity networks (RSN spatial maps) for use as targets in SBCA, using group-level independent component analysis (ICA; as implemented in FSL MELODIC; Beckmann and Smith, 2004; Beckmann et al. 2005; see also *Supplementary Materials and Methods*). This group-ICA resulted in the automatic estimation of 32 independent components in our HCs FMRI dataset.

Based on criteria and findings outlined previously (i.e., networks located largely outside of primary sensory cortical areas, with spatial characteristics reflecting neural systems implicated in dopamine-dependent processes, impulsivity or motivated behaviour; see also *Supplementary Materials and Methods* and Cole et al. 2013), eight RSNs of interest were selected for higher-level analyses. These eight (Supplementary Fig. 1) included: (i) a basal ganglia/limbic network (BGLN) comprising regions of bilateral striatum, pallidum, thalamus, midbrain and ventromedial/orbital prefrontal cortex (Fig. 1A, peak MNI coordinates in the left anterior putamen: x = −24, y = 6, z = 2), (ii) a fronto-temporo-parietal cortical ‘salience/executive’ network (SEN; Fig. 1B, left postcentral gyrus/central operculum peak: x = −64, y = −22, z = 16), (iii) right-lateralised (Supplementary Fig. 1C, right inferior parietal/superior occipital cortex peak: 46, −58, 48) and (iv) left-lateralised (Supplementary Fig. 1D, left lateral orbitofrontal/frontopolar cortex: −46, 44, −12) frontoparietal networks (FPNs), (v) anterior (Supplementary Fig. 1E, dorsomedial prefrontal cortex peak: 0, 46, 30) and (vi) posterior (Supplementary Fig. 1F, left precuneus/posterior cingulate cortex: −4, −54, 16) ‘default mode’ network (DMN) sub-systems, (vii) a cerebellar RSN (Fig. 1C, medial cerebellum peak: −2, −56, −30) and (viii) a somatomotor RSN (Supplementary Fig. 1H, right central sulcus/superior parietal cortex: 38, −26, 58).

**Figure 1.**
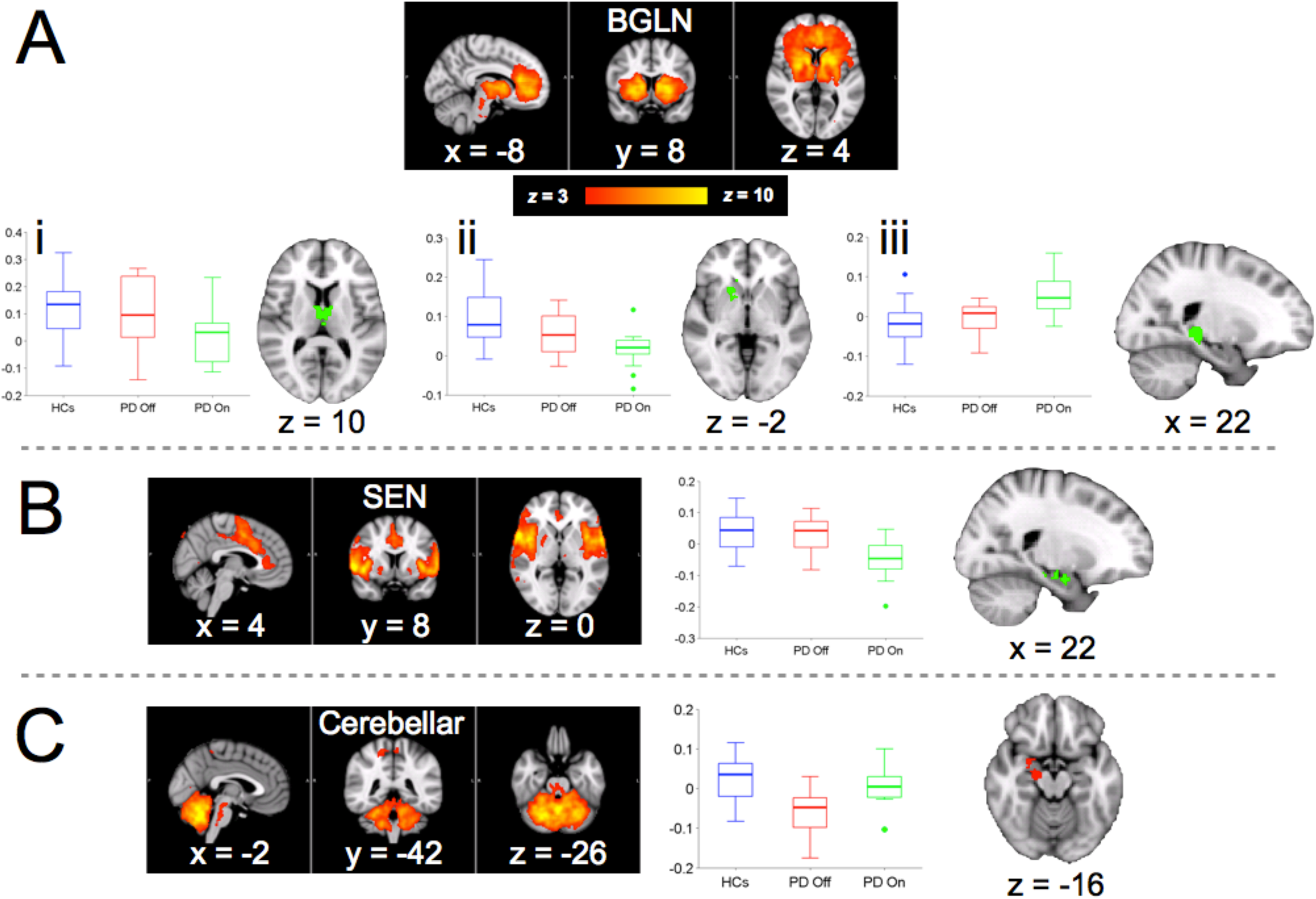
(A-B) Significant ‘non-normalising’ effects of DA treatment (i.e., normalising effects of DA *withdrawal*) on RSN cortico-subcortical functional connectivity measures (t > 2.3, p < 0.05, FWE-corrected), defined by SBCA (N = 34). (A) The BGLN displays reduced connectivity in PD On relative to PD Off and HCs with (i) bilateral anterior thalamus and (ii) right ventral-mid striatum/anterior pallidum, while displaying (iii) *increased* connectivity in PD On relative to PD Off and HCs in right posterior hippocampus. (B) A fronto-temporo-parietal SEN shows reduced connectivity with amygdala/anterior hippocampus in PD On relative to PD Off and HCs. (C) A significant ‘normalising’ effect of DA treatment (i.e., a non-normalising effect of DA *withdrawal*) on cerebro-cerebellar RSN functional connectivity measures (t > 2.3, p < 0.05, FWE-corrected). Connectivity between the cerebellar RSN and right amygdala, extending into midbrain regions is reduced in PD Off relative to PD On and HCs. Clusters of significant regional effects are shown in green (for non-normalising DA effects) or red (for normalising effects) on representative images. Red-yellow overlays depict regions of high functional connectivity within the RSNs themselves, as defined by HCs group-ICA. Boxplots display RSN-ROI correlation coefficients for each group and condition. Three-dimensional slice coordinates relative to the anterior commissure are provided in MNI152 stereotactic space (Montreal Neurological Institute, Canada). In all Figures, brain images are presented in line with radiological convention (left = right).

### Subject-wise seed-based correlation analysis (SBCA) of RSN connectivity with subcortical regions

Subject/session-specific measures of voxel-wise subcortical connectivity with each of the eight RSNs of interest (plus three ‘nuisance’ RSNs; see *Supplementary Materials and Methods*) were obtained using SBCA (see O’Reilly et al. 2010; Cole et al. 2013), for all HCs and PD patients in both treatment conditions separately. SBCA was carried out within a subject-specific, anatomically-derived subcortical seed mask. Every voxel within each ‘individualized’ seed mask was tested quantitatively in terms of its connectivity with each of the 11 RSN target maps as described previously (Cole et al. 2013; see also *Supplementary Materials and Methods*).

### Group-level analyses

The output of SBCA consisted of distinct subcortical spatial maps containing voxel-wise functional connectivity (correlation coefficient) values for each of the eight RSNs of interest, separately for each subject/condition (HCs, PD On and PD Off). Statistical contrasts in higher-level multiple regression analyses specifically examined RSN functional connectivity patterns for: (i) primary effects of DA treatment withdrawal in PD patients in a basic paired design (i.e., PD On >/< PD Off); (ii) ‘normalising’ effects of DA in PD patients (measured using contrasts comparing the connectivity in PD Off conditions to the average of that in HCs and PD On; i.e., 0.5 × HCs + 0.5 × PD On >/< 1.0 × PD Off); (iii) ‘non-normalising’ effects of DA in PD patients (0.5 × HCs + 0.5 × PD Off >/< 1.0 × PD On), the results of which could afford novel neurobiological interpretations in the context of frequently reported side effects of DA on certain cognitive functions in PD (e.g., Dagher and Robbins, 2009); and finally (iv) gross (treatment-independent) differences between HCs and PD patients (1.0 × HCs >/< 0.5 × PD On + 0.5 × PD Off). Significant regional effects were defined using nonparametric permutation testing (as implemented in FSL ‘randomise’) with cluster-mass thresholding (t > 2.3, p < 0.05) and full correction for family-wise error (FWE).

To investigate interactions between disease- or DA treatment-dependent connectivity patterns and impulsive decision-making as measured by the ITC questionnaire, bivariate within-group/condition correlations (Pearson’s r, p < 0.05, two-tailed) were carried out between individual subject measures of RSN-subcortical connectivity and mean delay discount rates (*k*-values) from the ITC scores (the latter were log-transformed to impose linearity prior to parametric testing). Significant clusters identified from the post-SBCA higher-level analysis were thus used as masks to extract mean connectivity scores from normalised (Fisher z-transformed) versions of RSN-specific correlation maps initially used as inputs to higher-level analyses. Statistical tests examining associations between impulsivity and these functional connectivity measures focussed on RSN connectivity relationships involving the ventral striatum that were affected by PD or DA treatment. To rule out any potential confounding effects of subject age and duration of treatment on ITC mean *k*-values in PD, bivariate correlations between these measures were also carried out.

## Results

### Non-normalising DA effects on cortico-subcortical RSN functional connectivity in PD

Within the current cohort of PD patients – and analogously to the results of their previously reported ITC behavioural data (Milenkova et al. 2011) – negative findings of basic DA medication withdrawal main effects were apparent when tested in isolation (see *Supplementary Results*). However, when exploiting the statistical power from additional quadratic contrasts including HCs, we identified a number of significant non-normalising effects of DA medication on cortico-subcortical RSN functional connectivity (cluster t > 2.3, p < 0.05, FWE-corrected). First, we found bilateral antero-medial thalamic functional connectivity with the BGLN to be reduced significantly in PD On relative to both PD Off and HCs (peak t = 5.46; coordinates: x = −6, y = −4, z = 12; Fig. 1A*i*). Second, a similar non-normalising DA connectivity effect was apparent between the BGLN and regions of the right (ventral and anterior mid-) striatum/putamen and anterior pallidum (t = 4.41; x = 20, y = 18, z = −4; Fig. 1A*ii*). Third, right posterior hippocampal/thalamic-BGLN connectivity (t = 4.22; x = 22, y = −36, z = −2) showed an ‘inverse’ non-normalising effect (PD On > HCs + PD Off; Fig. 1A*iii*). Finally, right anterior hippocampal and amygdalar regions (t = 5.11; x = 16, y = −14, z = −20) also showed a non-normalising effect of DA, in terms of reduced connectivity with the fronto-temporo-parietal SEN in PD On (Fig. 1B).

### Normalising DA effects on RSN functional connectivity in PD

A normalising effect of DA medication on connectivity (HCs + PD On > PD Off) was found between the cerebellar RSN and bilateral clusters located primarily within right (peak t = 4.72; x = 18, y = −16, z = −14) and left (t = 4.72; x = −18, y = −2, z = −22) amygdala, extending from/into midbrain regions (Fig. 1C). In addition, a normalising DA connectivity effect was apparent between the BGLN and regions of the left (ventral and anterior mid-) striatum/putamen and anterior pallidum (t = 4.34; x = −10, y = 4, z = −2). A graphical summary of normalising and non-normalising RSN connectivity effects of DA in PD is provided in Fig. 2. Additional significant, potentially ‘treatment-independent’ effects of disease on RSN cortico-subcortical connectivity are presented in Fig. 3 and described in detail in *Supplementary Results*.

**Figure 2.**
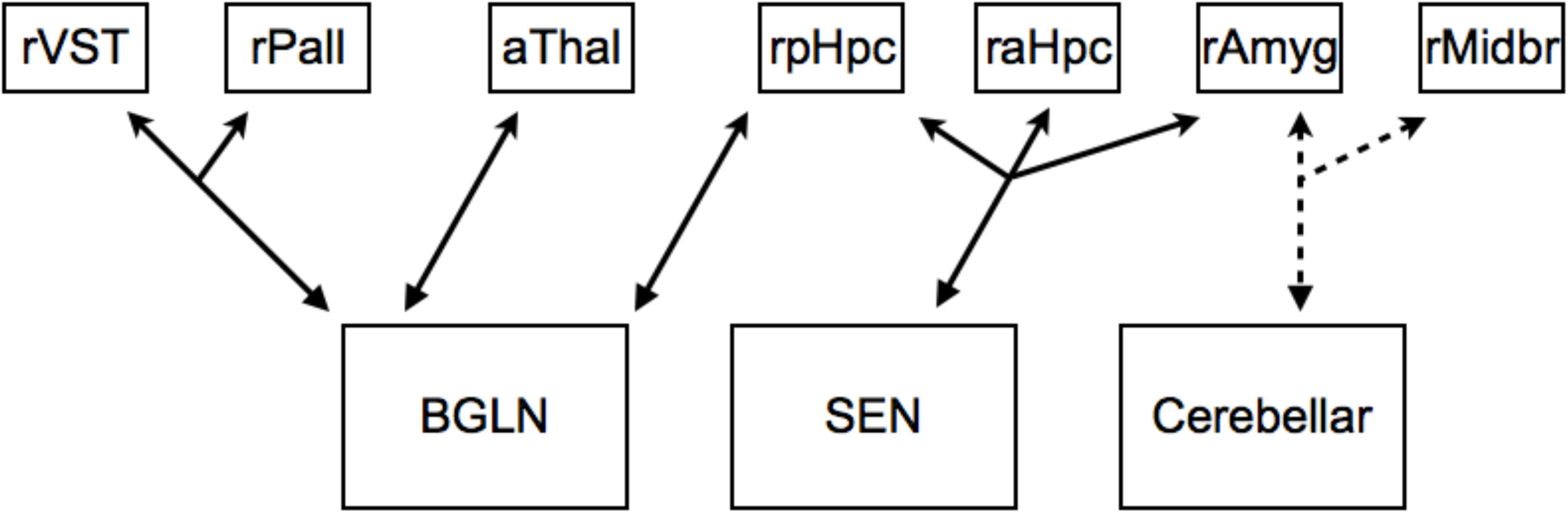
Schematic summary of significant non-normalised (solid arrows) and normalised (dashed arrows) DA-related functional connectivity relationships between distinct RSNs (bottom row) and subcortical regions (top row). ‘Split’ arrows indicate multiple anatomical regions that formed part of a single cluster identified by our higher-level analyses of DA-related RSN functional connectivity. rVST = right ventral striatum/putamen; rPall = right pallidum; aThal = bilateral anterior thalamus; rpHpc = right posterior hippocampus; raHpc = right anterior hippocampus; rAmyg = right amygdala; rMidbr = right midbrain.

**Figure 3.**
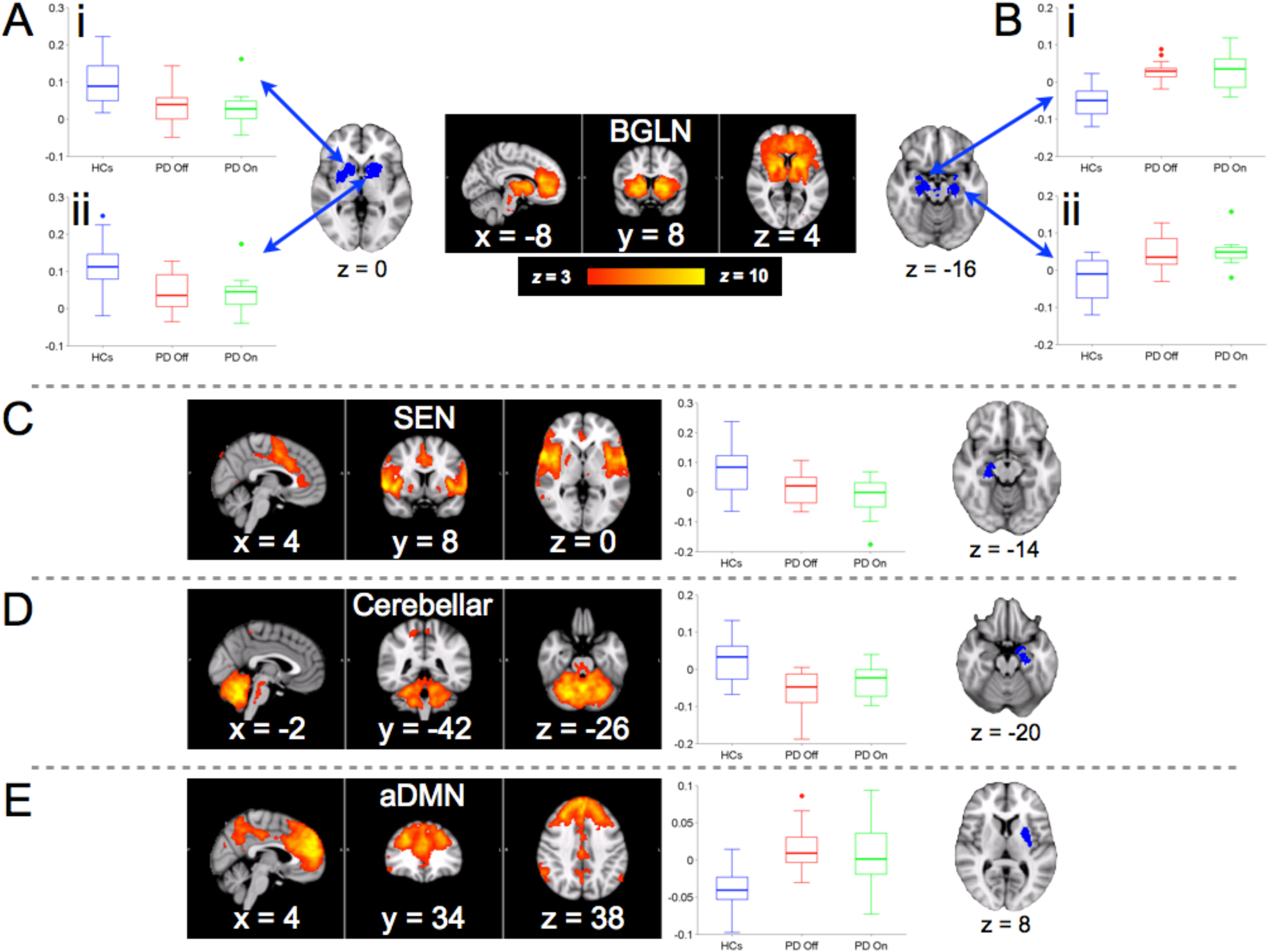
Significant, apparent ‘treatment-independent’, effects of PD on RSN cortico-subcortical functional connectivity measures (t > 2.3, p < 0.05, FWE-corrected), defined by SBCA (N = 34). (A) The BGLN displays reduced ‘within-network’ connectivity in bilateral (Ai, right; Aii, left) regions of striatum, pallidum and thalamus in PD patients relative to HCs. (B) BGLN connectivity with bilateral (Bi, right; Bii, left) amygdalo-hippocampal regions is increased in PD relative to HCs. (C) Right posterior hippocampus shows reduced connectivity with the fronto-temporo-parietal SEN in PD relative to HCs. (D) Reduced left amygdalo-hippocampal connectivity with the cerebellar RSN in PD. (E) Increased connectivity between left dorsal putamen and the anterior DMN in PD. Formatting as described in Fig. 1 legend.

### Association between PD-related, but not DA-related, connectivity modulations and impulsivity

Post-hoc correlations revealed a significant association in HCs (r = 0.48, p = 0.038), but in neither of the PD treatment conditions (p > 0.05), between ITC questionnaire measures of impulsive decision-making (log-transformed mean *k*-values) and BGLN connectivity within an extended striatal cluster identified in the image analysis (Fig. 4). This association was found specifically in the right hemisphere region (Fig. 3A*i*) displaying significantly greater BGLN connectivity in HCs relative to PD (independent of DA condition). Furthermore, this positive correlation was significantly different (p = 0.037) from an equivalent negative (but non-significant; r = −0.27) correlation in PD On. No significant correlations were found between ITC mean *k*-values and subject age or duration of treatment in PD, thus confirming a lack of influence of these factors on ITC-related RSN connectivity results.

**Figure 4.**
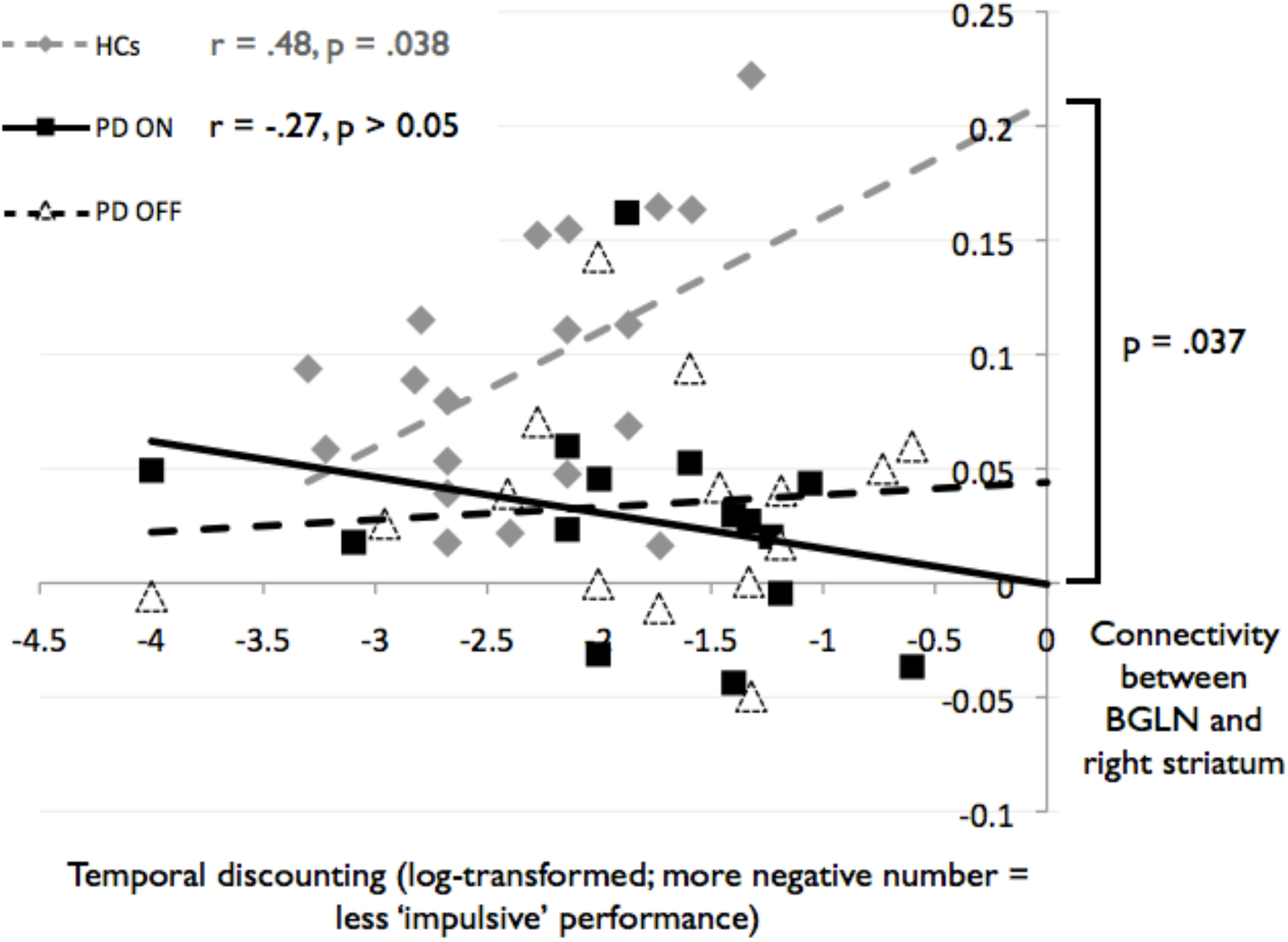
Association between BGLN-striatal functional connectivity and impulsivity in decision-making. A positive correlation (blue) was found between delay discounting behaviour (ITC questionnaire mean *k*-values) and BGLN-striatal connectivity in HCs (r = 0.48, p < 0.05, two-tailed; N = 19), but not in PD patients (red = PD On; green = PD Off; N = 15). This significant impulsivity-connectivity association was significantly different (p < 0.05, two-tailed) from the equivalent negative (but non-significant) correlation in PD On DA medications (but not during DA withdrawal).

## Discussion

In this study, we identified cortico-subcortical resting-state network functional connectivity patterns suggestive of treatment-dependent effects on systems-level processing in Parkinson’s disease relative to age-matched healthy control subjects. Specifically, by studying the effects of dopamine agonists in PD patients without clear neuropsychiatric comorbidities, considerable evidence was revealed for ‘non-normalising’ effects of DA on a range of cortico-subcortical functional connections (Fig. 2), echoing neuropsychological observations reported widely in the literature (Cools, 2006; Dagher and Robbins, 2009; Cilia and van Eimeren, 2011). Notably, in this study sample, individual differences in impulsive decision-making were found to be associated with connectivity within striatal/basal ganglia circuitry in healthy subjects, but not in the PD patients in whom this same circuitry displayed aberrant functional connectivity. Findings (i) support the existence of compensatory neural connectivity mechanisms in PD, via increased recruitment of cerebellar and other large-scale resting-state networks in response to striatal dysfunction (e.g., Appel-Cresswell et al. 2010; Helmich et al. 2012), while (ii) extending neurobiological evidence for the prior reported exacerbating effects on cognitive and motivational symptoms of modulating (increasing) dopamine neurotransmission pharmacologically, with commonly-prescribed DA drugs (e.g., Dagher and Robbins, 2009; Weintraub et al. 2010).

### Functional relevance of DA withdrawal effects on connectivity in PD

Previous studies in PD patients have highlighted both that (i) motor cortico-subcortical connectivity changes appear secondary to dopamine deficiency (Wu et al. 2009) and (ii) a key compensatory role exists for signalling involving cerebellar circuitry in mediating motor symptoms (Wu et al. 2011; Helmich et al. 2012). Indeed, here cerebellar RSN connectivity with the amygdala (and midbrain) *was* found to be normalised by DA drugs in patients. Future combined exploration of a broader range of neurochemical processes and whole-brain systems-level phenomena (e.g., cortico-subcortical, cerebro-cerebellar *and* cortico-cortical connectivity relationships) in PD may therefore be warranted. Having said this, the network most sensitive in this study to the effects of PD and DA on connectivity (in terms of the largest number of significant results identified) appeared to be the BGLN system, which comprises striato-thalamo-orbitofrontal circuitry implicated in dopamine function, reward processing, motivated behaviour and related neuropsychiatric disorders (Honey et al. 2003; Everitt and Robbins, 2005; Ersche et al. 2010; Koob and Volkow, 2010; Sesack and Grace, 2010). It is, therefore, likely to be one of the most relevant RSNs for studying brain functions and dysfunctions underpinned by dopamine and dopaminergic pathology in PD.

Incidences of addictive behaviours and impulse control disorders, such as compulsive gambling and hypersexuality, are reported with increasing regularity in PD patients, having been observed in up to 10% of those treated with DA drugs (Dagher and Robbins, 2009; Weintraub et al. 2010; Cilia and van Eimeren, 2011; Gao and Wu, 2016; Vargas and Cardoso, 2018). It is thus conceivable that cortico-subcortical networks subserving cognitive and motivational processes relevant for these behaviours would be altered differentially in PD On conditions, relative to PD Off and HCs (i.e., non-normalised). This assertion is supported by the current finding of differential associations, across groups and conditions, between large-scale cortico-subcortical network connectivity and impulsivity in decision-making. The inclusion criteria for this study dictated that no subjects, including PD patients, had comorbid addictions or other impulse control disorders. The absence of a direct correlation between ITC scores and connectivity in either treatment condition may, therefore, reflect this lack of overt behavioural pathology. However, a recent publication examining behavioural ITC data from subjects included in the current FMRI study found that, on average, DA withdrawal had no effect on delay discounting in PD, which was nonetheless increased overall in the patients relative to HCs (Milenkova et al. 2011). This finding of greater mean *k*-values in these PD patients could thus be indicative of group differences in impulse control at the sub-syndromal level, independent of short-term DA withdrawal effects. Nevertheless, even in the absence of diagnosed comorbid disorders, RSN connectivity metrics do appear to differentiate patient and control groups in terms of their within-group associations (or lack thereof, in the case of PD) with individual variability in impulsive choice. In short, findings imply that ‘healthy’ associations between impulsive decision-making tendencies and BGLN connectivity are disrupted in PD patients undergoing DA treatment.

### Compensatory neural mechanisms in PD

Both increases and decreases in RSN cortico-subcortical functional connectivity were found in PD patients, relative to age-matched HCs. It has been suggested previously that both increases in impulsive decision-making and neuroplastic reductions in striatal connectivity, which may be promoted by DA medications in PD, result from impaired dopaminergic functioning in basal ganglia (including striatum; e.g., Riba et al. 2008; Appel-Cresswell et al. 2010; Cai et al. 2011; Milenkova et al. 2011). Impulsive decision-making could therefore be a behavioural indication reflecting underlying neural mechanistic attempts to compensate for reduced striatal signalling in PD. This is corroborated by the current differential connectivity associations with individual differences in delay discounting (as measured by the ITC questionnaire), which showed more impulsive behavioural decision-making (i.e., a higher *k*-value) to be associated with significantly greater intra-BGLN connectivity in HCs, but with *reduced* (albeit not significantly so) intra-BGLN connectivity in PD On. In addition, it has been proposed that striatal motor dysfunctions in PD may be actively compensated for by increased functional recruitment of the cerebellum (Appel-Cresswell et al. 2010; Helmich et al. 2012; Wu and Hallett, 2013). Support for this theory is provided by the current observation of DA-normalised (reduced negative) cerebellar RSN connectivity with the right amygdala, although no clear association was discovered between the functional connectivity of this circuitry and motor function. Moreover, similar (but bilateral) amygdalar and hippocampal regions were found to display greater connectivity with the BGLN in PD relative to HCs; an effect of disease opposite in direction to the reduced striatal connectivity found within the BGLN. The current results thus also imply that compensatory functional circuitry in PD may extend beyond the cerebellum to more distributed cortical and subcortical networks.

### Methodological considerations

In this study, the repeated measures design within PD patients was not counterbalanced, as measurements were always taken firstly before, then during, DA medication withdrawal. This sequence was not only necessary for patient compliance, but also allowed the testing of clinically relevant treatment hypotheses related to ‘normalisation’ and ‘non-normalisation’ of functional connectivity by DA in PD, by maintaining direct experimental control of the timeframe of short-term medication withdrawal. It is conceivable that such a design feature could induce order effects that were difficult to account for in this study. This possibility could be explored, for example, by employing a placebo-controlled, crossover design in early stage, pre-medicated PD patients. However, symptomatology of cognitive or neuropsychiatric interest would be even less severe in such patients, thus any increase in sensitivity to the network-level functional connectivity effects of DA on the underlying neurocircuitry would be unlikely. Future studies investigating the systems-level effects of DA *per se* may, therefore, benefit from examining populations with related, but less debilitating, DA-treated disorders in which more prolonged drug withdrawal periods would be possible, such as restless legs syndrome.

A second important methodological issue for the current study design was the variability in treatment type and regimen across PD patients (see *Materials and methods*). Specifically, some patients took DA monotherapy, while others were taking concomitant L-dopa medication. The latter was not withdrawn from those patients. Furthermore, doses of DA and L-dopa, as well as the exact DA medications being taken, varied across patients. No controlling measures could easily be applied here for these potential sources of variability, as it was important to limit interference with patients’ ongoing medical care to the minimum required to test the study hypotheses. This constraint, along with the procedures limiting the DA withdrawal period to approximately 12 hrs, could provide an explanation for why only a single normalising effect of DA on connectivity was found in the current sample. However, the fact that many non-normalising DA effects *were* identified, even in PD patients with no comorbid behavioural disorders of inhibition or addiction, implies that such confounds do not, comparatively, influence sensitivity of the systems-level metrics applied here to the primary experimental pharmacological manipulation. Nonetheless, future investigations might benefit from alternative (e.g., cross-sectional) designs enabling direct assessment of the influences of these multiple neuropharmacological factors on large-scale network connectivity patterns relevant for PD and other disorders of neurochemistry.

Finally, it is necessary to highlight the potential for functional connectivity differences between subjects or groups to be influenced by structural anatomical variability within neuronal networks. This is particularly pertinent to research conducted in neurological populations, who may exhibit varying degrees of macro- or micro-structural pathology in the underlying grey or white matter that account for some of the variability in BOLD signal fluctuation synchronicity (e.g., Voets et al. 2012). The current study controlled for the possible (if unclear) impact of this in PD, by employing an ‘inclusive’ method of subcortical seed definition in subject-level SBCA and restricting group-level analyses to subcortical space defined by PD patients (see *Supplementary Materials and Methods*). These steps were taken in order to avoid biasing results towards being influenced excessively by gross structural differences between groups or sub-groups of individuals. Nonetheless, as noted (and relative to Alzheimer’s disease, for example), comparatively little agreement and clarity exist currently regarding the contribution to functional deficits, or indeed the existence, of structural neuropathology in non-demented PD patients (Burton et al. 2004; Dalaker et al. 2009; Menke et al. 2010; Jubault et al. 2011; Tinaz et al. 2011), such as those sampled in the current study.

### Conclusions

This study has characterised diverse systems-level functional connectivity modulations associated with Parkinson’s disease and dopamine agonist treatments. Our findings provide brain systems-level evidence to support current theories that: (i) while DA drugs are known to improve motor symptoms of nigrostriatal dopamine degeneration, they may also promote unwelcome cognitive changes, such as increased impulsivity, in predisposed individuals through hyperstimulation of the mesolimbic dopamine pathway; and (ii) neural mechanisms compensating for dysfunctional striatal and basal ganglia signalling patterns in PD proceed via increased recruitment of large-scale cerebro-cerebellar and cortico-subcortical systems. Overall, the findings identify functional connectivity pathways that may prove useful targets for future neuromodulatory intervention studies testing, for example, treatment efficacy or side effects in PD and other disorders in which individual neurochemical predispositions influence therapeutic and behavioural outcomes.

## Acknowledgements

The authors would like to thank the patients who participated in this research. The authors are grateful to Dr. Natalie Voets for providing helpful comments on a manuscript draft.

