## Supplementary Materials for "Effects of dopamine agonist treatment on resting-state network connectivity in Parkinson’s disease"

**Supplementary Materials and Methods**

*Participant details*

All subjects were drawn from a sample described, along with other non-imaging procedures, in a previously published article (Milenkova et al. 2011). Patients had idiopathic Parkinson's disease (PD), as diagnosed by an experienced neurologist, without comorbid depression, dementia or impulse control disorders (i.e., no history of gambling problems, compulsive shopping, hypersexuality, punding, or abuse of their medication) and were judged to be able to function to a reasonable degree (e.g., they did not express pronounced motor symptoms) when their medication was withdrawn for approximately 12 hrs prior to scanning. Healthy control subjects (HCs) were all free from neuropsychiatric disorders.

*Intertemporal choice (ITC) questionnaire*

The ITC questionnaire provides a measure of 'delay discounting' of monetary rewards for individual subjects, which refers to a reduction in the subjective value of a larger later reward, relative to that of a smaller earlier one, as the temporal delay to the former increases (Kirby et al. 1999). Delay discounting in ITC has been shown to occur with increased frequency in disorders associated with maladaptive reward processing, including addictions, pathological gambling and impulse control disorders in PD (Kirby et al. 1999; Petry, 2001; Housden et al. 2010; Voon et al. 2010; Milenkova et al. 2011). However, relationships between this measure of impulsivity, brain dopamine function and individual differences in large-scale cortico-subcortical network functional connectivity remain less clear, as do the specific identities of the key connectivity pathways mediating these associations (e.g., Marco-Pallares et al. 2010; Peters and Büchel, 2010; Camchong et al. 2011; Kayser et al. 2012).

The ITC questionnaire was administered at each visit as described in detail in a recent publication examining behavioural data from subjects also included in the current functional magnetic resonance imaging (FMRI) study (Milenkova et al. 2011). In brief, the ITC requires subjects to choose in sequence between paired monetary rewards, one smaller and immediate and the other larger and delayed. Subjects were incentivised to make realistic decisions with the information that they would later have the opportunity to actually achieve one of these reward choices through a probabilistic die-throw. A measure, *k*, of the delay discount rate was produced for each subject. The *k*-value computation follows a (quasi-) hyperbolic function such that a higher value for *k* indicates that the participant in question subjectively devalued delayed rewards at a progressively steeper rate than an individual displaying a lower *k*-value. A version of the ITC questionnaire sensitive to the effects of different reward magnitudes on delay discounting was employed, although as the original behavioural study found no evidence of differential magnitude effects between groups and conditions (Milenkova et al. 2011), the mean *k*-value is the variable of primary interest in the current context.

*Subcortical seed mask definition*

We created anatomically-defined seed masks of the entire subcortex for each subject, for use in subject-level seed-based partial correlation analysis (SBCA; O'Reilly et al. 2010). The caudate, putamen, ventral striatum, globus pallidus, amygdala, hippocampus and thalamus were segmented automatically from each subject's T1 structural image using FSL FIRST (Patenaude et al. 2011) and combined into a single mask image for each subject. As FIRST does not segment midbrain regions containing the brain's primary dopaminergic output neurons, the substantia nigra and the ventral tegmental area (e.g., Everitt and Robbins, 2005; Sesack and Grace, 2010), a midbrain mask covering these regions and consisting of six binary, bilateral volumes was extracted from the Talairach Daemon atlas (Lancaster et al. 2000; labels = midbrain, substantia nigra, subthalamic nucleus, red nucleus, mammillary body and medial geniculum body). Nonlinear warp transformation of this atlas-derived midbrain mask to the high-resolution space of each subject was then carried out using FSL FNIRT. Finally, these subject-specific combined seed masks were affine registered to echo planar imaging (EPI) space (using FSL FLIRT) for subsequent subject-wise SBCA.

The subcortical mask used for higher-level analyses in MNI standard space was formed from the subject-level masks of PD patients *only*. The aim of this was to provide a comparatively conservative masking approach, to better control for the potential (although inconclusively established) brain structural differences between groups impacting on observations of functional processes (e.g., Burton et al. 2004; Dalaker et al. 2009; Menke et al. 2010).

*Group-ICA and network definition details*

The resting-state FMRI data of HCs were thus entered into probabilistic multi-session independent component analysis (ICA) with temporal concatenation (as implemented in FSL MELODIC; Beckmann and Smith, 2004; Beckmann et al. 2005). This group-ICA approach decomposed the concatenated 4-D dataset (174 volumes per scan × 19 subjects = 3306 image volumes) into spatial maps of structured signal in the data (and associated time courses), identifying components, including resting-state networks (RSNs), displaying consistent spatiotemporal coherence within scans and maximal spatial independence across subjects. The number of components for the HCs dataset was estimated automatically using the Laplace approximation to the Bayesian evidence for the model order in a probabilistic principal component model (for details see Beckmann and Smith, 2004). This group-ICA resulted in the automatic estimation of 32 independent components. Twenty-one of these were deemed to represent artefacts of motion, non-neuronal physiology or magnetic susceptibility (see e.g., Beckmann and Smith, 2004; Beckmann et al. 2005) and thus they were excluded from further analyses. Based on their spatial characteristics, the remaining 11 components were judged to correspond to known RSNs. Three of these represented primary sensory brain systems (located in visual cortical regions) for which we did not hypothesise connectivity modulations in PD or by dopamine agonist (DA) treatments. Of the remaining eight RSNs, six included frontal cortical regions reported to mediate impulsivity in decision-making, to be sensitive to mesolimbic/mesocortical dopaminergic neuromodulation and thus hypothesised to display non-normalising effects of DA on functional signalling (Honey et al. 2003; Kelly et al. 2009; Koob and Volkow, 2010; Cools and D'Esposito, 2011; Cole et al. 2012a, 2012b), while the seventh and eighth comprised motor cortical and cerebellar regions potentially relevant for the therapeutic (normalising) functional effects of drug modulation of nigrostriatal dopamine in PD.

Notably, the spatial characteristics of the BGLN described in the group-ICA results from the current control dataset are not confined purely to subcortical regions. Although the 'peak' basal ganglia/limbic network (BGLN) voxel is highly proximal, for example, to that of the equivalent RSN identified by our group previously (Cole et al. 2012b), in the current study the BGLN also demonstrated large nodes of high functional connectivity within extended portions of ventromedial and orbital prefrontal cortex (Fig. 1A). This system strongly matches circuitry described previously as "large-scale striato-thalamo-orbitofrontal networks implicated in the processing of natural rewards and the regulation of behavior" (Ersche et al. 2010). Furthermore, it comprises regions known to be modulated by dopamine and compromised in addiction and other disorders associated with impulsive/compulsive behaviours (Honey et al. 2003; Everitt and Robbins, 2005; Koob and Volkow, 2010; Sesack and Grace, 2010). We thus included the BGLN as a network of interest in the current post-SBCA analysis. Similarly, the cerebellar RSN (Fig. 1C) was included as emerging evidence has highlighted a compensatory role for cerebellar functioning and connectivity in PD, which is hypothesised to arise in response to impaired dopaminergic function in striatal regions and their reduced integration with large-scale networks (Appel-Cresswell et al. 2010; Helmich et al. 2012). Finally, the somatomotor RSN was included because DA medications are used widely to treat motor system dysfunction in PD.

Prior to conducting subject-wise SBCA, we employed two procedures to control for the potential (but relatively unclear) contribution of tissue macro- or micro-structural variations to regional neuropathology in non-demented PD patients (Burton et al. 2004; Dalaker et al. 2009; Menke et al. 2010; Jubault et al. 2011; Tinaz et al. 2011). Firstly, to ensure more accurate alignment of neural systems defined at a group level to the brain space of individuals, prior to affine transformation to EPI space, RSN target maps from the group-ICA of HCs were *nonlinearly* (as opposed to linearly) transformed from MNI152 space to the high-resolution space of each subject (as implemented in FSL FNIRT); wherein, secondly and prior to registration to EPI space, voxels failing to exceed a liberal threshold of > 10% probability of containing grey matter (as calculated using FSL FAST) in the equivalent T1 structural were excluded from these 'subject-specific' RSN spatial maps. These procedures were applied to the data of all subjects, whether PD patients or HCs.

*SBCA of RSN connectivity with subcortical regions – details*

In subject-specific SBCA, subcortical seed masks were examined individually, in EPI space, for their voxel-wise spatial distributions of functional connectivity strength with the characteristic activity of each of the 11 RSNs. Voxel-wise connectivity strengths were quantified by calculating partial correlation coefficients between the BOLD signal time series at each mask voxel and that of the weighted principal eigenvariate associated with each RSN (the latter calculated via subject-wise principal component analyses; see O'Reilly et al. 2010). Voxel-wise coefficients are termed 'partial' because the analysis associated with a given target RSN controlled for (i.e., regressed out) the seed voxel's activity relationship with each of the other 10 RSNs examined as targets in separate subject-wise correlation analyses. All 11 non-artefactual components from group-ICA were thus included in these first-level analyses to ensure that potential extraneous interactions, or temporally overlapping relationships, between any of the eight RSNs of interest (see main manuscript *Higher level analyses*; Supplementary Fig. 1A-H) or with any of the three nuisance RSNs (networks located chiefly in visual cortical areas) could be factored out of the analysis, in effect treating the latter as confound regressors.

In these analyses we also controlled for the confounding influences of structured noise from white matter (WM) and cerebrospinal fluid (CSF) tissue types and residual motion artefacts. To this end, binary T1-segmented maps of WM and CSF (calculated using FAST) were registered to EPI space using FLIRT and, for each session, used as masks against the associated, preprocessed functional datasets, in order to extract confound time series that were calculated as the mean BOLD signal within these tissue masks. As well as these WM and CSF confounds, six time series resulting from the motion correction procedure (as implemented in FSL MCFLIRT) describing individual subject translational and rotational head motion parameters were temporally filtered in the same manner as the FMRI data and also regressed out of the SBCA. Mean absolute and relative head motion did not differ significantly between DA treatment sessions or subject groups (all $p > 0.39$, two-tailed t-tests).

### Supplementary Results

*PD On vs. PD Off: no basic effect of dopamine agonist medication withdrawal on RSN functional connectivity*

Within the current population of PD patients, paired analyses revealed no significant effects of DA medication on cortico-subcortical RSN functional connectivity (cluster $t < 2.3$, $p > 0.05$, FWE-corrected). This finding of negative basic medication effects in paired analyses of PD patients' FMRI data mirrors that of their ITC behavioural data (Milenkova et al. 2011). One possible explanation for this sparseness of main effects is that, due to the lengthy biological half-lives of most DA medications, the withdrawal period of 12 hrs used here might limit the sensitivity of this simple within-subject contrast (see also main article *Discussion*). We therefore hypothesised further that comparing PD On and Off treatment conditions differentially relative to un-medicated HCs, using statistical contrasts probing 'normalising' or 'non-normalising' effects, might provide increased sensitivity to variability across conditions and reveal additional DA effects on higher-order RSN functional connectivity.

*Apparent 'treatment-independent' cortico-subcortical RSN functional connectivity differences in PD*

In addition to the described effects of DA medication and modifications therein (Fig. 2, main article), a number of (seemingly treatment-independent) disease-driven effects on cortico-subcortical RSN functional connectivity were identified. A significant reduction in BGLN functional connectivity was found between bilateral clusters in (ventral and extended) striatum/putamen and pallidum, extending into subthalamic and midbrain nuclei and anterior thalamus (right: peak $t = 5.26$; $x = 18$, $y = 14$, $z = 0$; left: $t = 5.57$; $x = -10$, $y = 4$, $z = 0$) in PD relative to HCs (Fig. 3A). This widespread basal ganglia regional effect overlapped with the (more spatially constrained) region of right ventral/anterior mid-striatum described above as displaying a non-normalising connectivity effect of DA (as well as the regions showing a similar, but more extensive, thalamic effect; see first *Results* section and Fig. 1A*i-ii*, main article). Conversely, bilateral hippocampal and amygdalar regions (extending into midbrain and right posterior thalamus; right t = 5.99; x = 24, y = -28, z = -12; left t = 4.56; x = -18, y = -20, z = -16) showed *increased* treatment-independent BGLN connectivity in PD relative to HCs (Fig. 3B). In the right hemisphere, there was some overlap between these regions and the posterior hippocampus cluster found to display a non-normalising BGLN connectivity increase with DA (Fig. 1A*iii*, main article). Additional gross effects of disease were found, including reduced left hippocampal-amygdalar connectivity with the cerebellar RSN (t = 4.00; x = -24, y = -26, z = -16; Fig. 3D) and increased left dorsal putamen connectivity with the anterior DMN in PD relative to HCs (t = 4.12; x = -34, y = -8, z = 6; Fig. 3E). Finally, a treatment-independent effect was observed of PD reducing the connectivity between the somatomotor RSN and bilateral antero-medial thalamus, extending into the left dorsal posterior putamen and subthalamic nucleus (right t = 3.92; x = 8, y = -10, z = 8; left t = 3.75; x = -4, y = -18, z = 12).

Fig S1

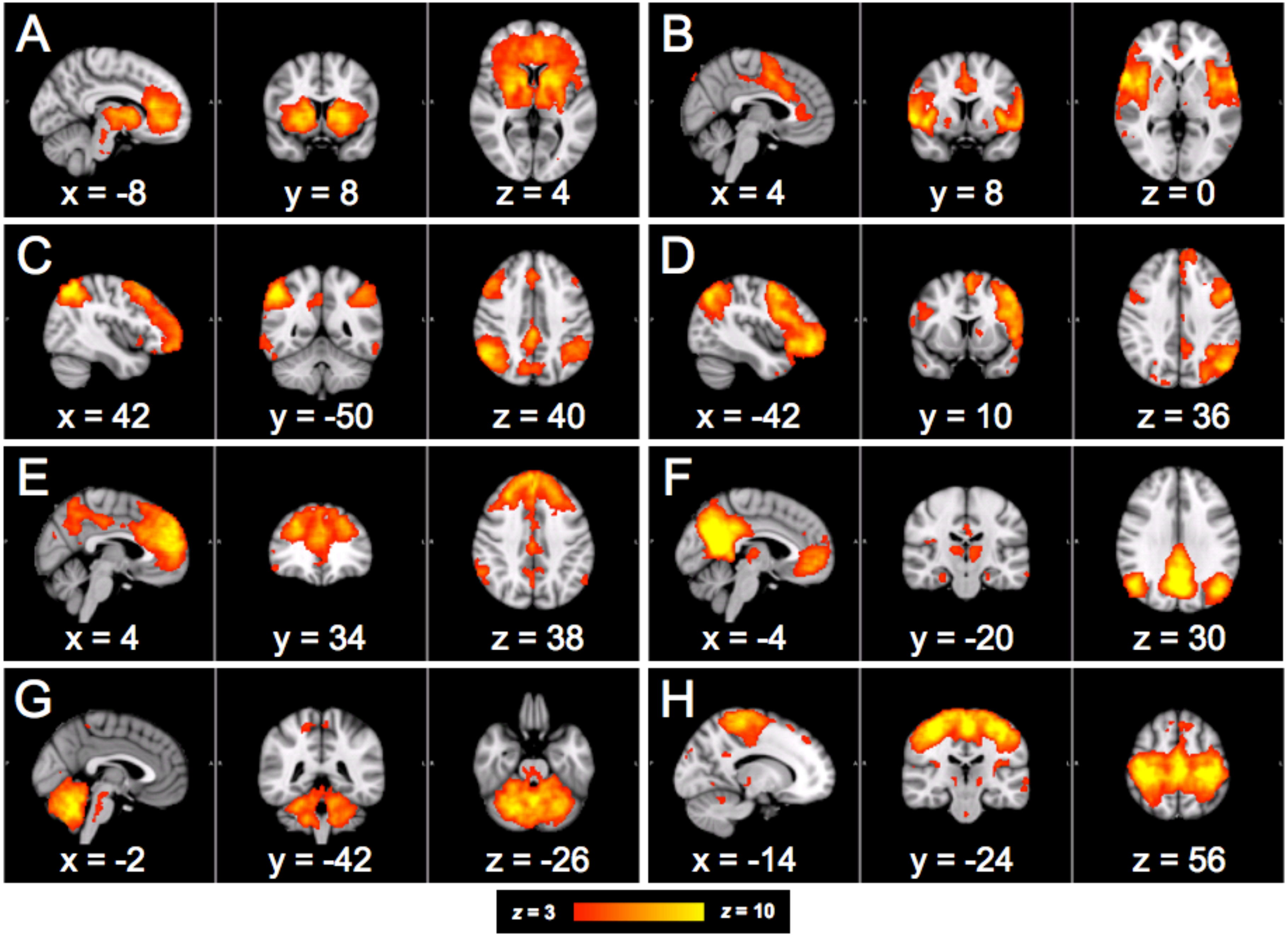
A
x = -8
y = 8
z = 4
B
x = 4
y = 8
z = 0
C
x = 42
y = -50
z = 40
D
x = -42
y = 10
z = 36
E
x = 4
y = 34
z = 38
F
x = -4
y = -20
z = 30
G
x = -2
y = -42
z = -26
H
x = -14
y = -24
z = 56
z = 3
z = 10
